# Most-probable number based minimum duration of killing assay for determining the spectrum of rifampicin susceptibility in clinical *M. tuberculosis* isolates

**DOI:** 10.1101/2020.06.29.179200

**Authors:** Srinivasan Vijay, Hoang Ngoc Nhung, Nguyen Le Hoai Bao, Do Dang Anh Thu, Le Pham Tien Trieu, Nguyen Hoan Phu, Guy E. Thwaites, Babak Javid, Nguyen T. T. Thuong

## Abstract

Accurate antibiotic susceptibility testing is essential for successful tuberculosis treatment. Recent studies have highlighted the limitations of minimum inhibitory concentrations (MIC) based phenotypic susceptibility methods in detecting other aspects of antibiotic susceptibilities in bacteria. Duration and peak of antibiotic exposure, at or above the MIC required for killing the bacterial population, has emerged as another important factor for determining the antibiotic susceptibility. This is broadly defined as antibiotic tolerance. Antibiotic tolerance can further facilitate the emergence of antibiotic resistance. Currently there are limited methods to quantify antibiotic tolerance among clinical *M. tuberculosis* isolates. In this study, we develop a most-probable number (MPN) based minimum duration of killing (MDK) assay to quantify the spectrum of *M. tuberculosis* rifampicin susceptibility within subpopulations, based on time duration of rifampicin exposure required for killing the bacterial population. MDK_90_-_99_ and MDK_99.99_ defined as the minimum time duration of antibiotic exposure at or above MIC required for killing 90-99% and 99.99% of the initial (pre-treatment) bacterial population respectively. Results from the rifampicin MDK assay applied to 28 laboratory and clinical *M. tuberculosis* isolates showed that there is variation in rifampicin susceptibility among isolates. Rifampicin MDK_99**/**99.99_ time for isolates varied from less than 2 to 10 days. MDK duration was correlated with larger sub-populations of *M. tuberculosis* from clinical isolates that were rifampicin tolerant. Our study demonstrates the utility of MDK assays to measure the variation in antibiotic tolerance among clinical *M. tuberculosis* isolates and further expands clinically important aspects of antibiotic susceptibility testing.

## Introduction

Tuberculosis (TB), caused by *Mycobacterium tuberculosis*, results in more than 1.5 million deaths a year (1). Although TB can be successfully treated with antibiotics, a minimum six months of treatment is required (2). Relapse post-treatment and emergence of antibiotic resistance are major negative sequelae of inadequate treatment (2). Success in therapy requires accurate determination of antibiotic susceptibility, and initiation and adherence to an effective antibiotic regimen (3).

The most common measure of antibiotic susceptibility of *M. tuberculosis* isolates is the minimum inhibitory concentration (MIC) (4), the concentration of antibiotics that inhibits, kills or reduces the growth of at least 99% of the bacterial population (5). The MICs for antibiotics vary between clinical *M. tuberculosis* isolates, and epidemiologically relevant MIC break-points are used to define *M. tuberculosis* isolates as susceptible or resistant for each antibiotic (6).

However, it has recently emerged that MICs in the susceptible range, below the clinical cut-off for resistance, can be associated with poor treatment outcomes (7). But resistance to an antibiotic, defined by the MIC, is not the only measure of antimicrobial susceptibility. Strains with identical MICs can still display widely different susceptibilities to bactericidal antibiotics, a phenomenon described as antibiotic tolerance (8, 9). With antibiotic tolerance, subpopulations of bacteria that are genetically susceptible to an antibiotic are killed more slowly than the bulk population (9). Studies have identified emergence of mutations in *M. tuberculosis* clinical isolates associated with increased levels of antibiotic tolerance (10-13). Emergence of antibiotic tolerance can further facilitate the emergence of resistance (14-17). These studies highlight limitations of reliance on MIC alone as the only measure of antibiotic susceptibility, and the possible role of antibiotic tolerance in poor treatment progression and evolution of antibiotic resistance in *M. tuberculosis* isolates. Hence, it is important to investigate the level of antibiotic tolerance among clinical *M. tuberculosis* isolates.

One way to measure the level of antibiotic tolerance is to determine the time duration of antibiotic exposure at MIC or above MIC concentrations required for killing majority of the bacterial population (18). Kill-curve assays allow study of the dynamics of bacterial population killing post antibiotic exposure (19). Antibiotic-mediated killing of bacterial populations is bi-phasic, an initial rapid killing of the majority population followed by slow-rate of killing of minority populations, also known as tolerant and persistent bacterial populations (19). Repeated antibiotic exposure can lead to an increase in the level of antibiotic tolerance both at sub-population or bulk-population level (11, 20). This will further increase the time duration required for killing the bacterial population. Recent clinical studies have shown in-host evolution of antibiotic tolerance in other pathogenic bacteria and its association with treatment complications (21).

Therefore, quantitative assays to measure antibiotic tolerance of bacterial populations are required. Commonly used tolerance assays are based on the minimum time duration required for killing the bacterial population by antibiotics, also known as minimum duration of killing assay (MDK) (18). The MDK assay can be used to measure both bulk-population and sub-population levels of antibiotic tolerance (18). MDK_90_ and MDK_99_ refer to the minimum time duration required for killing one or two log-fold (90 to 99%), or majority of the bacterial population during antibiotic exposure. Hence, MDK_90_ or MDK_99_ can quantify the bulk-population level of antibiotic tolerance (18). Whereas, MDK_99.99_ determines the minimum time duration required for killing 4 logs of the population and, assuming the tolerant sub-population is > 0.1%, quantifies the sub-population level of antibiotic persistence (18).

Reliable and feasible methods are required to measure MDK values of clinical *M. tuberculosis* isolates, to determine if there are significant variations in antibiotic tolerance among *M. tuberculosis* isolates. A critical measure used in the MDK assay is to accurately determine the viable bacterial number at different time points post antibiotic exposure of the bacterial culture. Usually, bacterial viability is measured by either colony forming units (CFU) or most-probable number (MPN) methods (22). In the MPN method, the viable bacterial numbers in culture is determined by generating 10-fold serial dilutions of the culture until limiting dilution (i.e. the dilutions show no visible bacterial growth over time) is reached. The viable bacterial number (MPN/mL) is calculated by considering that the highest dilution showing growth contains at least one viable bacterium and then multiplying by the dilution factor (23). Previous studies have shown that MPN and CFU methods give comparable viable counts for *M. tuberculosis* (24) and the MPN method further facilitates *M. tuberculosis* growth from clinical samples (25), since in the MPN method all serial dilutions can be simultaneously incubated in one microtiter plate, overcoming the laborious process of plating multiple serial-dilutions to obtain countable numbers of colonies using CFU method (26).

Given the prime importance of rifampicin in drug-sensitive tuberculosis treatment (27), we developed and tested an MPN-based MDK assay for determining the level of rifampicin tolerance among *M. tuberculosis* isolates. We applied this assay on a pilot scale to determine the spectrum of rifampicin susceptibility in 26 clinical *M. tuberculosis* isolates and identified variation in rifampicin susceptibility among these isolates, which included 6 isoniazid and rifampicin susceptible, 19 isoniazid resistant and 1 MDR-TB, resistant to both isoniazid and rifampicin.

## Materials and methods

### Bacterial isolates

*M. tuberculosis* H37Rv, *M. bovis*-BCG from laboratory strain collection and clinical *M. tuberculosis* isolates from pulmonary tuberculosis patients, collected during pre and post treatment for a previous study (28) were revived from archive and initially cultured in 7H9G medium (supplemented with 10% OADC, 0.2% glycerol). Followed by one or two subculture in Middlebrook 7H9T medium (BD Difco™, Thermo Fisher Scientific, USA) supplemented with OADC, 0.05% tween 80, with or without 0.2% glycerol, these cultures were used for the MDK experiments.

### Mycobacterium culture for MDK assay

Mycobacterial isolates were cultured in 10 – 15 ml 7H9T medium in 50 mL tubes, with or without glass beads, in shaking incubator at 37°C. When the culture reached optical density (O.D_600_) range of 0.4-0.6, acid fast staining was done to confirm the purity of the *Mycobacterium* culture. Culture set with glass beads was vortexed for 3 minutes to disrupt bacterial aggregates, as the aggregates may influence the level of antibiotic tolerance (29). Cultures in both set of tubes, with or without glass beads, were diluted to O.D_600_ 0.4 with fresh 7H9T medium and used for the MDK assay.

### Rifampicin preparation

Rifampicin (Sigma-Aldrich, USA) stock solutions were prepared in DMSO filter sterilized and aliquots were stored at −20°C for MDK assay.

### Bacterial viability determination by most-probable number method

for MDK assay, first initial viability of mid-log (O.D_600_ 0.4) *M. tuberculosis* cultures were measured by MPN method (0 day), by taking 1 mL aliquot from the 0.4 O.D culture and harvesting the culture at 7000 RPM for 5 minutes; bacterial cell pellet was washed once by suspending in 1 mL fresh 7H9T medium and re-centrifuged, then re-suspended in 1 mL 7H9T medium. 100 µL from this 1 mL culture was transferred to 96 well plates as undiluted culture in triplicate for serial dilution. 10 µL from the undiluted (10°) culture was used for serial dilution in 90 µL 7H9T media up to 10^−9^ dilutions in microtiter plates (Figure 1). Each time new pipette tips were used for mixing and transferring culture during serial dilution. Final two wells, with 7H9T media at the end of serial dilution after 10^−9^, were left for sterility check without adding any mycobacterial culture (Figure 1). The serially diluted plates sealed by 96 well plastic adhesive seal and incubated at 37°C, without shaking for one or two months. Immediately after initial (0 day) viability measurement, the mycobacterial cultures in the 50 mL tubes were treated with rifampicin at a final concentration of 1 or 2 µg/mL and cultures were further incubated at 37°C with shaking (Figure 1). At different time points (1, 2, 3, 5, 10, 15 and 20 days, depending on experiments) post-rifampicin treatment, viable bacterial number were measured again by removing 1 mL from the rifampicin treated culture, cells were washed once with fresh 7H9T medium to remove rifampicin and the MPN assay was repeated as performed on the initial day (0 day) before rifampicin treatment (Figure 1).

**Figure 1.**
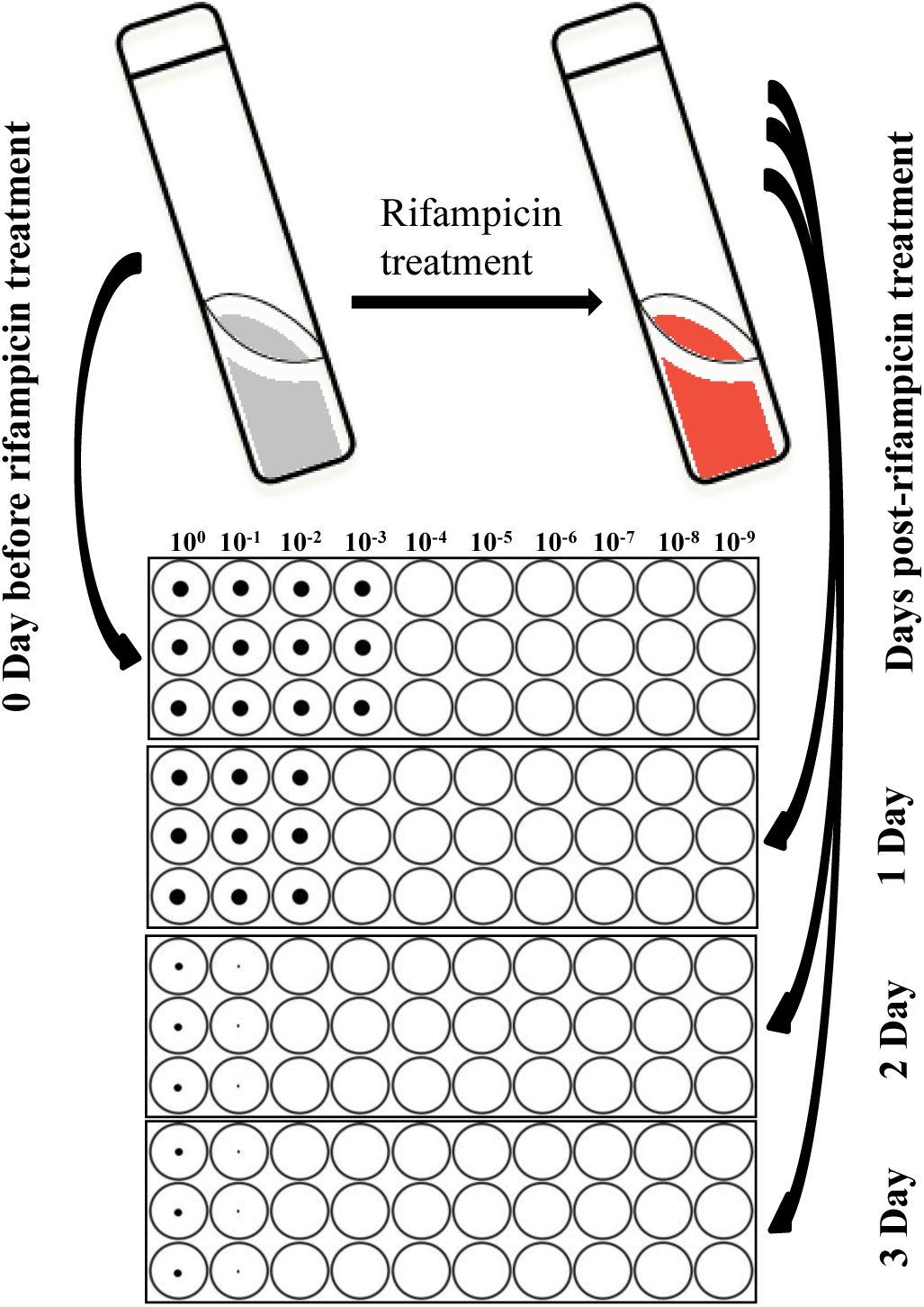
MDK assay study design. Mycobacterial culture were grown in 50 mL tube until O.D reached 0.4-0.6, this culture was diluted to O.D 0.4 with fresh 7H9T medium. 1 mL from this culture was removed at 0 day (just before rifampicin treatment), for measuring viable bacterial number by serial dilution in triplicate using 96 well microtiter plates. The serially diluted microtiter plates were incubated for 15-20 days for determining the MPN/mL. After 0 Day sampling the remaining culture in the tube was treated with rifampicin (1 or 2 µg/mL) and incubated further. At different time points post-rifampicin treatment, 1 mL rifampicin treated culture is removed, centrifuged, cell pellet is washed free of antibiotic, re-suspended in fresh media and serially diluted similar to 0 day. The viable bacterial number in the original culture before and after rifampicin treatment is determined by the MPN method.

Growth in 7H9T medium at the bottom of 96 well plates were recorded by the Thermo Fisher Sensititre Vizion digital MIC viewing system (Thermo Fisher, Scientific Inc., USA) after 15-20 days, one and two months of incubation (Figure 1 and Fig. S1). Final MPN results were calculated based on 15-20 days of incubation for antibiotic tolerance determination between clinical *M. tuberculosis* isolates considering different factors influencing the MPN number (Fig. S1). There was some increase in MPN at one month (possibly due to post-antibiotic effect or resuscitation of non-replicating persisters) and drying up of well was also observed in 96 well plates at one and two month time point (Fig. S1). Early reading at 15-20 days clearly distinguished difference in the extent of growth even at same serial dilution between *M. tuberculosis* isolates (Fig. S1A). Due to higher sensitivity in detecting difference in growth at 15-20 days incubation and drying up of wells at later incubation period, we used 15-20 days MPN results for calculating survival fraction post-rifampicin treatment (Fig. S1). *M. tuberculosis* viability at each time point is calculated as mean MPN/mL with 95% CI (confidence interval) or with standard deviation. For single set of dilution series, the MPN/mL was determined by the highest dilution showing growth, multiplied by the dilution factor. Based on MPN value of each time point, survival curve were plotted to determine the killing dynamics post-rifampicin treatment. The MDK_99_ and MDK_99.99_ time were calculated based on the time duration required for 99% and 99.99% reduction in the survival of the population respectively during the killing phase, as compared to the initial MPN/mL taken as 100%.

### Drug susceptibility testing

Mycobacterial Growth Indicator Tube (MGIT) was used for phenotypic drug susceptibility testing (DST) by BACTEC MGIT 960 SIRE Kit (Becton Dickinson) according to the manufacturer’s protocol. BACTEC MGIT 960 SIRE DST was done for streptomycin (1.0 µg/mL), isoniazid (0.1 µg/mL), rifampicin (1.0 µg/mL) and ethambutol (5.0 µg/mL) (28).

### GeneXpert MTB/RIF

For the GeneXpert MTB/RIF detection of rifampicin resistance, we followed the manufacture protocol using the GeneXpert instrument, GeneXpert Dx system software and cartridge (Cepheid) (30). Briefly, 0.2 mL of decontaminated sputum was added into 2 mL of sample reagent and transferred into a test cartridge. The cartridge was inserted into the test platform of a GeneXpert instrument and post-run analyzed for detection of *M. tuberculosis* and rifampicin resistance (30).

## Results

### Developing an MPN-based MDK assay for determining *M. tuberculosis* rifampicin susceptibility

In order to develop the MDK assay and to determine rifampicin susceptibility, we initially optimized the assay by testing laboratory strains *M. tuberculosis*-H37Rv and *M. bovis*-BCG treated with rifampicin in mid-log phase (Figure 2A, B). Both laboratory strains showed rapid killing with 1 µg/mL rifampicin treatment, the cut-off concentration used for defining susceptible and resistant isolates in Mycobacterial Growth Indicator Tube (MGIT) culture for clinical isolates (31). The decrease in MPN/mL compared with day 0 in microtiter plates showed a MDK_99_ time of less than 1 day for *M. bovis*-BCG and between 2 and 3 days post-rifampicin treatment for H37Rv with and without bead beating respectively (Figure 2B). Indicating additional step of bead beating to remove bacterial aggregation did not show any difference in H37Rv rifampicin tolerance compared to cultures without beads but only with tween 80 for disrupting the bacterial aggregates.

**Figure 2.**
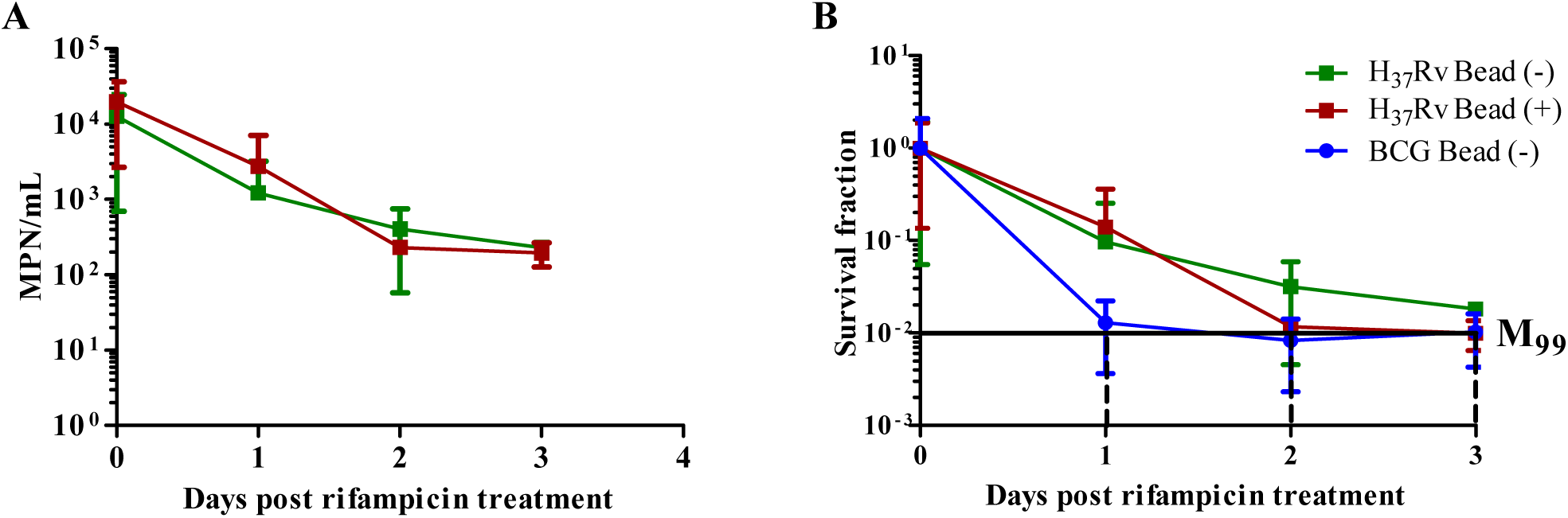
MPN based MDK assay development using laboratory mycobacterial strains *M. tuberculosis* H37Rv and *M. bovis* BCG. (A) Mid-log cultures of laboratory strain H37Rv (with and without beads) treated with rifampicin 1 µg/mL and viable bacterial number measured as MPN/mL just before rifampicin treatment (0 day) and each day post-rifampicin treatment for three days. (B) Survival fraction and MDK_99_ of *M. tuberculosis* H37Rv and *M. bovis* BCG, black solid horizontal line indicate two log-fold reduction in survival fraction (M_99_) as compared to 0 day. Dashed vertical line indicate the MDK_99_ time for individual isolate with or without beads. The data is average MPN/mL at each day for 4 experiments. *M. tuberculosis* H37Rv with and without beads are marked with different colors.

### Applying rifampicin MDK assay for clinical *M. tuberculosis* isolates

We then applied the MDK assay to investigate variation in rifampicin susceptibility among clinical *M. tuberculosis* isolates (n = 26). Clinical *M. tuberculosis* isolates were selected based on GeneXpert MTB/RIF results for rifampicin susceptibility and phenotypic DST by MGIT. For this assay, we included laboratory strain H37Rv (Rv), 6 isoniazid and rifampicin susceptible isolates (Figure 3A, B), 19 isoniazid resistant but rifampicin susceptible isolates (Figure 3C) and a MDR-TB isolate (R), resistant to both isoniazid and rifampicin (Figure 3 A-C). For the MDK assay in clinical isolates, we used higher rifampicin concentration of 2 µg/mL for both H37Rv and clinical *M. tuberculosis* isolates cultured without bead beating and survival fraction was determined by the MPN method using a single set of serial dilution at different time points (Figure 3). The increase in antibiotic concentration increased the time duration of maintaining rifampicin concentration above the clinical susceptibility cut-off of 1 µg/mL in the medium. This extended the duration of the killing phase, as well as the range of MDK time for determining the rifampicin susceptibility among clinical *M. tuberculosis* isolates. Laboratory strain H37Rv and MDR-TB isolate were included as controls for high rifampicin susceptibility and resistance respectively.

**Figure 3.**
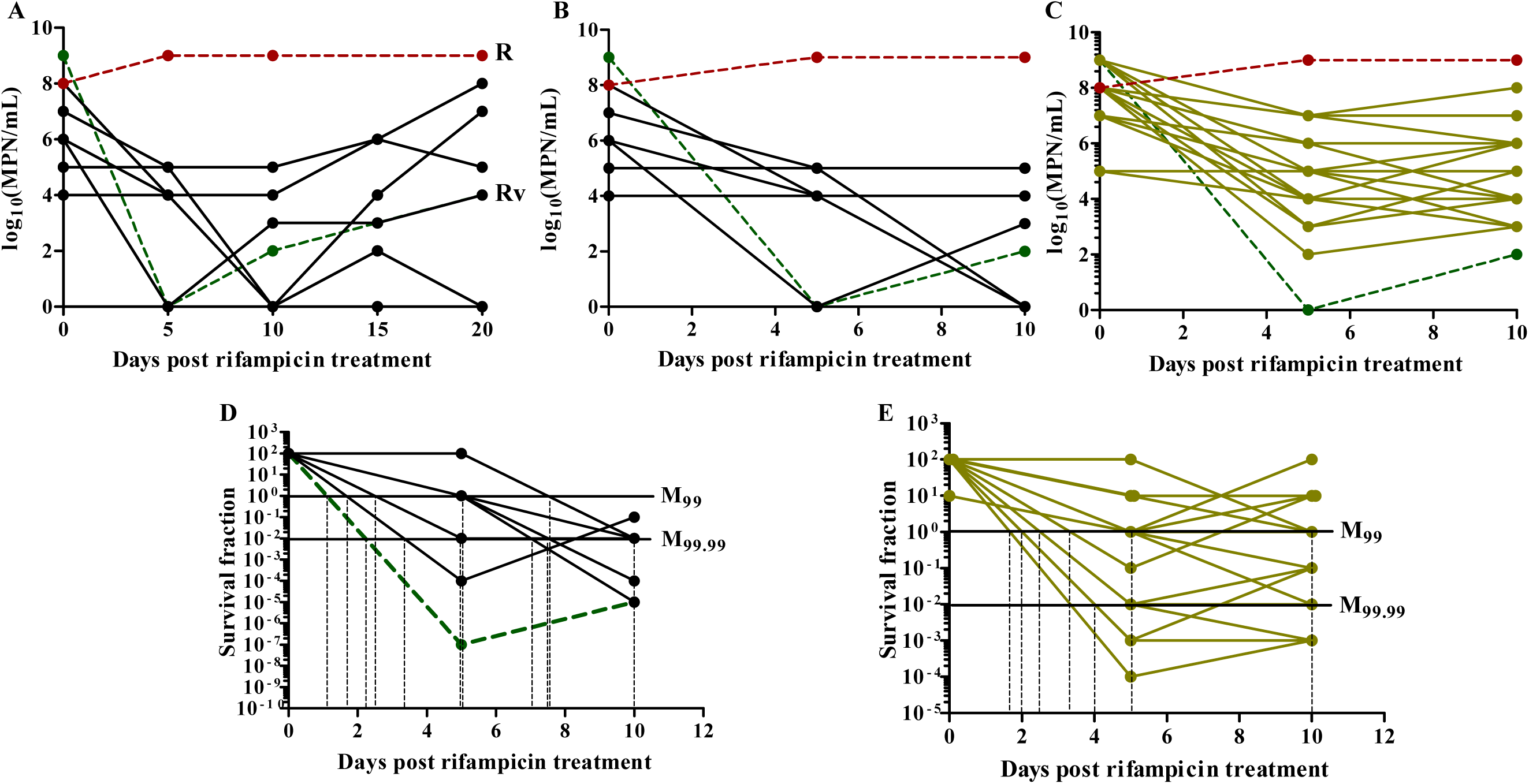
Applying rifampicin MDK assay to clinical *M. tuberculosis* isolates. (A) Mid-log cultures of laboratory strain H37Rv (Rv, green colored dash line), 6 clinical *M. tuberculosis* isolates, susceptible to isoniazid and rifampicin (black solid lines), and MDR-TB isolates (R red colored dashed line) were treated with 2 µg/mL of rifampicin. Viable mycobacterial cell number was determined by MPN method at 0 day (just before rifampicin treatment), 5, 10, 15 and 20 days post rifampicin treatment. (B) Mycobacterial survival was analyzed only up to 10 days post rifampicin treatment as regrowth of *M. tuberculosis* isolates were observed after 5-10 days of rifampicin treatment. (C) Rifampicin MDK assay for 19 isoniazid resistant (colored solid lines), H37Rv and MDR-TB (green and red colored dash lines) isolates treated initially with 2 µg/mL rifampicin. (D) normalized survival fraction of susceptible and (E) isoniazid resistant isolates to determine the MDK_99_ and MDK_99.99_ time, black solid horizontal lines indicate two log-fold (M_99_) and four log-fold (M_99.99_) reduction in survival fraction as compared to 0 day (normalized as 100% for susceptible and isoniazid resistant isolates). Black dashed vertical lines show approximate time required for 99% or 99.99% reduction in survival fraction of individual *M. tuberculosis* isolates.

All the rifampicin-susceptible *M. tuberculosis* isolates and H37Rv showed reduction in viability, i.e. killing phase for 5-10 days of treatment compared with day 0 MPN/mL, whereas rifampicin-resistant MDR-TB isolates showed growth or maintenance of MPN/mL compared with day 0 over 20 days of rifampicin treatment (Figure 3A). From 5 to10 days of rifampicin treatment re-growth phase was observed in most of the clinical *M. tuberculosis* isolates (Figure 3A), potentially representing outgrowth of *de novo* rifampicin resistant cells (15). Since MDK time can be measured only during the killing phase of rifampicin treatment (32), the range of MDK time measurements in our assay was restricted from 2 to 10 days of rifampicin treatment for susceptible (Figure 3B, D) and isoniazid resistant isolates (Figure 3 C, E). There was substantial variation in population and sub-population levels of rifampicin tolerance among susceptible and isoniazid resistant *M. tuberculosis* isolates as determined by approximate MDK_99_ and MDK_99.99_ time (Figure 3, D-E). H37Rv showed high rifampicin susceptibility, similar to the earlier observation with 1 µg/mL rifampicin treatment, having an MDK_99_ and MDK_99.99_ time of less than 2 and 3 days respectively, whereas for susceptible and isoniazid resistant clinical *M. tuberculosis* isolates MDK_99_ time distribution varied from 2 to > 10 days and MDK_99.99_ time varied from 3 to > 10 days, more than 10 days indicate high level of rifampicin tolerance beyond the detection limit of our assay (Figure 4). There was no statistical significant difference between overall rifampicin tolerance distribution, measured as MDK_99_ and MDK_99.99_ time, between susceptible and isoniazid resistant groups (Figure 4A, Mann-Whitney U). We used the median of MDK_99_ (5 days) and MDK_99.99_ (10 days) distribution to group susceptible and isoniazid resistant isolates with low (≤ median) and high (> median) rifampicin tolerance (Figure 4A, B), all high rifampicin tolerance isolates were beyond the 75^th^ percentile or above the upper detection limit of our assay (Figure 4A, B). Out of 19 isoniazid-resistant isolates, 5 and 10 isolates showed significantly high rifampicin tolerance as determined by MDK_99_ and MDK_99.99_ respectively compared to susceptible and other isoniazid resistant isolates (Figure 4A, B).

**Figure 4.**
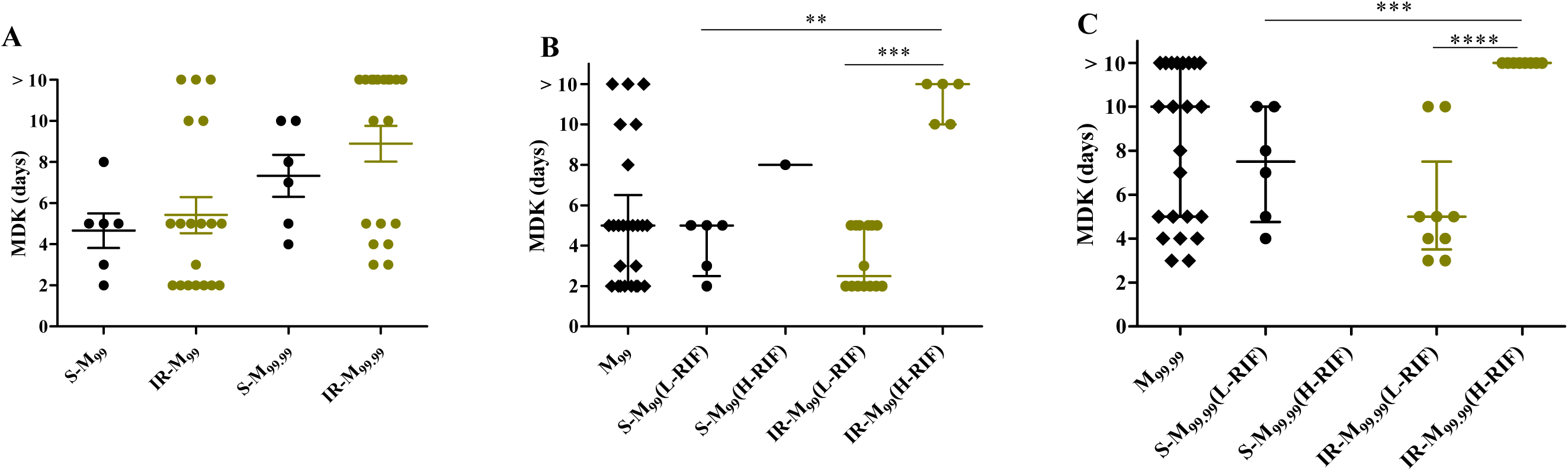
Variation in rifampicin tolerance as determined by MDK time. (A) MDK_99_ (M_99_) and MDK_99.99_ (M_99.99_) time for susceptible (S) and isoniazid resistant isolates (IR). (B) M_99_ and (C) M_99.99_ for all susceptible and isoniazid resistant isolates combined to determine the median (50th percentile) in the first column, median value for M_99_ = 5 days and M_99.99_ = 10 days were used to group susceptible and isoniazid resistant isolates with low (L-RIF ≤ median) and high (H-RIF > median) rifampicin tolerance. Each data point represent one clinical *M. tuberculosis* isolate, median and interquartile range of distributions are given (p – values represent ** < 0.01, *** < 0.001, and **** < 0.0001, Mann-Whitney U).

To further confirm the robustness and reliability of the MDK assay for clinical isolates, we repeated the assay with 3-6 independent biological replicates of 6 strains, representing fully drug-susceptible, isoniazid-resistant, and MDR-TB. The repeat assay fully recapitulated the observations, again revealing that 3/4 of the isoniazid-resistant isolates had showed longer MDK times for rifampicin compared with the fully drug-susceptible isolate (Fig. S2).

Our data confirms the ability of the MDK assay to discriminate differences in the bulk-population and sub-population susceptibility and tolerance to rifampicin among clinical *M. tuberculosis* isolates.

## Discussion

In this study we developed an MPN-based MDK assay for determining the spectrum of rifampicin susceptibility in clinical *M. tuberculosis* isolates. Our MDK assay can be easily adapted to study the variation in killing dynamics and determine the susceptibility level for different conditions such as different concentrations of rifampicin, other anti-tuberculosis antibiotics, combinations of antibiotics, and host stresses or bacterial metabolic adaptations (such as different carbon sources) by changing the treatments accordingly. Our pilot-scale study indicates the presence of bulk-population and sub-population level variation in rifampicin susceptibility among clinical *M. tuberculosis* isolates, as determined by MDK_99_ and MDK_99.99_ time respectively.

Rifampicin-susceptible clinical *M. tuberculosis* isolates (fully susceptible and isoniazid-resistant, n = 25) show variation in MDK time ranging from less than 2 up to 10 days, and greater than 10 days which is beyond the detection limit of our assay using rifampicin 2 µg/mL. This indicates the ability of MDK assay to distinguish a spectrum of rifampicin susceptibility, ranging from highly susceptible to highly tolerant.

Tolerance or susceptibility assays require reliable methods to determine the survival fractions of bacterial populations at different time durations post antibiotic exposure. Previous studies have shown that MPN method quantitatively works as good as, and sometimes even better than CFU method for determining the viability of *M. tuberculosis* (24, 25). In addition, it has an advantage of incubating all serial dilutions, which is essential for determining the antibiotic tolerance among clinical *M. tuberculosis* isolates with wide variations in the spectrum of tolerance. Adopting the MPN method of the MDK assay in 96-well microtiter plates significantly reduces the labor required for testing antibiotic tolerance in large numbers of clinical isolates as compared with the CFU method. Furthermore, the MPN method also reduces the time required for reading the results as presence or absence of growth at each dilution (visual check), compared with the time required for counting colonies in the CFU method.

A striking observation of our study was the relatively increased tolerance to rifampicin in isoniazid-resistant isolates, in particular with regards to MDK_99.99_. Routine laboratory MIC testing can miss such tolerance in clinical isolates^27^. Importantly, association of high rifampicin tolerance with isoniazid resistance may further contribute to the *de novo* emergence of multi-drug resistance or failure of multi-drug combinational therapy (17, 33) and require further detailed investigation. The spectrum and high level of antibiotic tolerance observed in our clinical isolates is in concordance with the in-vitro evolution of antibiotic tolerance in laboratory strain of mycobacterium (11) and high level of antibiotic tolerance observed in other bacteria (34).

This assay has been developed considering laboratories based in low and middle income countries and the growth in the microtiter plate can be read using a simple mirror box. The viability can be detected early using fluorescence dyes to further reduce the incubation time and obtain rapid results for clinical application (35). MDK assay will also help us to screen for phenotypic antibiotic tolerance among clinical *M. tuberculosis* isolates, and identify novel genetic variants and molecular mechanisms associated with the emergence of antibiotic tolerance (10-12). In addition, MDK assay can also help to identify possible chemical agents to reverse such tolerance by increasing the antibiotics potential (16). Investigating antibiotic tolerance and its emergence in clinical *M. tuberculosis* isolates will greatly improve treatment strategies by identifying hard-to-treat phenotype and stratifying treatment approach (36), prevent relapse or emergence of antibiotic resistance (17, 33).

## Acknowledgments

We thank Saurabh Mishra and Carl Nathan, Department of Microbiology and Immunology at Weill Cornell Medicine for providing MPN calculation program.

